# *Montastraea cavernosa* corallite structure demonstrates distinct morphotypes across shallow and mesophotic depth zones in the Gulf of Mexico

**DOI:** 10.1101/402636

**Authors:** Michael S. Studivan, Gillian Milstein, Joshua D. Voss

## Abstract

This study assessed morphological variation of the depth-generalist coral *Montastraea cavernosa* across shallow and mesophotic coral ecosystems in the Gulf of Mexico (GOM) using thirteen corallite metrics. While corallite structure differed significantly across sites, we observed that mean corallite diameters were smaller and spacing was greater in mesophotic corals as compared to shallow corals. Additional corallite variation, including greater mean corallite height of mesophotic samples, are hypothesized to be photoadaptive responses to low light environments. Multivariate analyses also revealed two distinct morphotypes identified by significant variation in corallite spacing with >90% accuracy. A ‘shallow’ morphotype was characterized by larger, more closely-spaced corallites, while a ‘depth-generalist’ type exhibited smaller, further-spaced corallites. Variable presence of morphotypes within some sites suggests genotypic influence on corallite morphology as there was a slight, but significant, impact of morphotype on genetic structure within shallow zones in the Flower Garden Banks. Patterns of increased algal symbiont (Symbiodiniaceae) density and chlorophyll concentration were retained in the depth-generalist morphotype even in shallow zones, identifying multiple photoadaptive strategies between morphotypes. The results of this study suggest that morphological variation among *M. cavernosa* represents a combination of genotypic variation and phenotypic plasticity rather than responses to environmental stimuli alone.

## Introduction

Using morphology as the sole method of species delineation can be confounded in scleractinian corals due to the considerable morphological variation observed within some species [1–6]. Occurrences of homologous morphological characteristics among coral species that are genetically distant from one another suggest further limitations for taxonomy based solely on morphology [7]. In one notable case in the Tropical Western Atlantic (TWA), high levels of skeletal variation in the *Orbicella* (previously *Montastraea) annularis* species complex, in conjunction with observed niche partitioning, resulted in the split of the complex into three sister species [8–10]. With additional genotyping efforts, *M. cavernosa* was determined to be the only species within the genus *Montastraea*, while the remaining three species formerly of the *M. annularis* species complex were reassigned to the genus *Orbicella* [5].

Further investigation of corallite variation and feeding behavior within *M. cavernosa* identified two morphotypes in Panama [11–14], leading to one theory of cryptic speciation within the species. A diurnal morphotype had smaller but continuously open polyps, while a nocturnal morphotype was characterized by larger polyps that were expanded only at night. There were also significant differences in algal symbiont density and colony respiration between the two morphotypes, where larger polyps allowed greater photosynthetic yields but subsequently higher respiration rates [11]. Currently, there is little molecular or reproductive evidence to support the claim of cryptic speciation among the morphotypes [15,16], but surveys along a depth gradient in Puerto Rico uncovered different vertical distributions for the morphotypes. The majority of shallower (6 m) colonies were of the smaller diurnal morphotype, while 60% of the deeper (20 m) colonies were the larger nocturnal morphotype, and the author proposed that morphological variation was due to different environmental conditions across depths and tradeoffs between photosynthesis and feeding rates [17]. It is important to note, however, that a study analyzing *M. cavernosa* morphology across the TWA observed two comparable morphotypes with no clear pattern across depths in Belize, implying that depth is not the sole factor affecting corallite phenotype [15]. Many other coral species have demonstrated remarkable levels of morphological variation across environmental gradients as well [18], but the underlying factors and mechanisms are relatively unknown.

Variation in skeletal structure among individuals is thought to be the interactive result of environmental stimuli and genotype. There is debate whether genotype contributes more heavily to coral morphology [19], or whether a combination of both genotype and environmental conditions act as simultaneous drivers [9,20,21]. Tests of environmental versus genotypic influence on phenotype through reciprocal transplant experiments have demonstrated that the environment has significant influence on coral morphology, but that genotype may limit the degree of morphological plasticity [22]. Previous studies have also identified a significant interaction between genotype and environmental conditions following transplantation, meaning genetic factors contributed to variation in skeletal morphology differently over a range of environmental conditions [23,24]. Differences in intracolony variation across populations provides further evidence for a genotypic influence on coral morphology [25], given the varying degrees of genotypic diversity observed across reefs at multiple spatial scales [26,27]. With the increasing number of genetic markers available for coral species, renewed investigations of variation across natural populations and with manipulative experiments are warranted to better understand the ability of coral individuals and populations to adapt to environmental variation across environmental gradients and multiple habitats.

Currently, there is limited understanding of intraspecific morphological variation across shallow and mesophotic coral ecosystems (∼30–150 m) [6,28,29], primarily due to a lack of morphological data collected beyond 20 m. Recent studies of *M. cavernosa* in Bermuda observed smaller corallites and overall colony sizes in mesophotic compared to shallow zones [30,31]. The authors suggested that micromorphological plasticity in this species was the result of photoadaptation and not necessarily selection as there was limited evidence of genetic differentiation between depth zones [32]. While there is a general understanding of environmental characteristics that may influence coral physiology and morphology in mesophotic zones (reviewed in [28,42,43]), environmental data including downwelling irradiance and zooplankton abundance are lacking for most mesophotic habitats including those in the Gulf of Mexico. Similar ecological processes that influence the genetic connectivity of coral populations across shallow and mesophotic coral ecosystems are only beginning to be explored. Connectivity of coral populations across depths is contingent on reproductive compatibility between shallow and mesophotic conspecifics and may be affected by potential morphological lineages within a species [11,15,17,31,35]. It is therefore important to assess whether morphological variation exists within coral populations and to determine whether that variation is reflective of genetic structure and connectivity. Depth-generalist species present an opportunity to observe phenotypic variation of a species within a small vertical scale to minimize any potential interactive effects of reef environments. Similarities in coral morphology across depth zones may not only indicate the possibility for morphotypes within species to adapt to different environments, but also reveal the presence of genetic influences on morphological plasticity.

Through examination of micromorphological variation in the depth-generalist species *M. cavernosa* across a wide range of geographical locations, we determined whether there exists a significant shift in corallite structure between shallow and mesophotic coral populations. Furthermore, we aimed to address whether evidence of adaptation to low-light environments is represented in algal symbiont and genotypic variation across populations, through comparison of morphological characteristics to symbiont and population genetics data from the same samples [36–38].

## Materials and methods

### Study sites and sample collection

Natural populations of *Montastraea cavernosa* were sampled to assess morphological variation among shallow and mesophotic sites in the northwest and southeast Gulf of Mexico (NW GOM and SE GOM, respectively). In total, 212 *M. cavernosa* samples were collected across six sites in the GOM during expeditions in 2014–2016 (S1 Fig). Colonies were photographed *in situ* and sampled haphazardly at least one meter apart to minimize the likelihood of sampling clones. Prior to sampling, colonies were observed briefly for any conspicuous patterns of polyp behavior. Coral fragments approximately 15–25 cm^2^ in area were collected from the margins of *M. cavernosa* colonies by SCUBA divers with hammer and chisel or by remotely-operated vehicle (ROV) with five-function manipulator and suction sampler. Coral samples were collected from Flower Garden Banks National Marine Sanctuary (FGBNMS) under permits FGBNMS-2010-005 and FGBNMS-2014-014. Vertically-contiguous reef habitats exist within the Flower Garden Banks (FGB), allowing direct comparison of morphology between shallow and mesophotic depth zones within sites. For these collections, relatively shallow samples were collected at 20 m on the coral caps of West and East FGB (WFGB and EFGB, respectively), and mesophotic samples were collected at 45 m along the bank margins. In total, 59 coral fragments were collected from West FGB (shallow, n=30; mesophotic, n=29) and 59 fragments from East FGB (shallow, n=37; mesophotic, n=22). Samples were collected at 50 m from the mesophotic-only habitats of Bright Bank (BRT, n=19) and McGrail Bank (MCG, n=26). In the SE GOM, 25 fragments were collected from the mesophotic-only Pulley Ridge (PRG) at 65 m and 24 fragments were collected from the nearby shallow Dry Tortugas (DRT) at 29 m. Fragments were processed with a dental water pick (Waterpik Water Flosser) to remove coral tissue and subsequently bleached in a 10% sodium hypochlorite solution to remove any remaining connective tissue or surface skeletal color.

### Morphological characters

Samples designated for morphometric analyses required five undamaged corallites and intact neighboring corallites; additionally all corallites measured were at least one row of corallites away from colony margins [25]. Thirteen morphometric characters were identified from previous studies of morphological variation in *M. cavernosa* [2,17,25,39]. All metrics were quantified for preliminary analysis on a subset of available samples (Dry Tortugas, 25–33 m, n=5) to determine if any characters could be eliminated while still maximizing morphological variation captured. Five of the characters lacked significant variation across samples, had strong correlations with other metrics, or included inherent variability that may have compromised the ability to recognize variation across samples (*e.g.* costal structures were frequently eroded in between corallites and therefore produced inconsistent length measurements across corallites; S1 and S2 Tables). Eight remaining morphometric characters were used in subsequent analyses (Table 1, Fig 1). Corallite and theca height were measured to the nearest 0.01 mm using dental calipers (ProDent USA). Scaled photographs were taken by a Canon G12 camera with a 6.1–30.5 mm lens (∼10–20 mm focal length) with the target corallite centered to minimize edge distortion and ensuring the corallite surface was perpendicular to the lens angle using a bubble level. The remaining metrics were measured using the scaled photographs in ImageJ [40,41]. All metrics were measured four times per corallite across five corallites, resulting in 20 replicate measurements per sample (Table 1), except in the case of corallite spacing where distance to all neighboring corallites were measured (Fig 1C).

**Fig 1.**
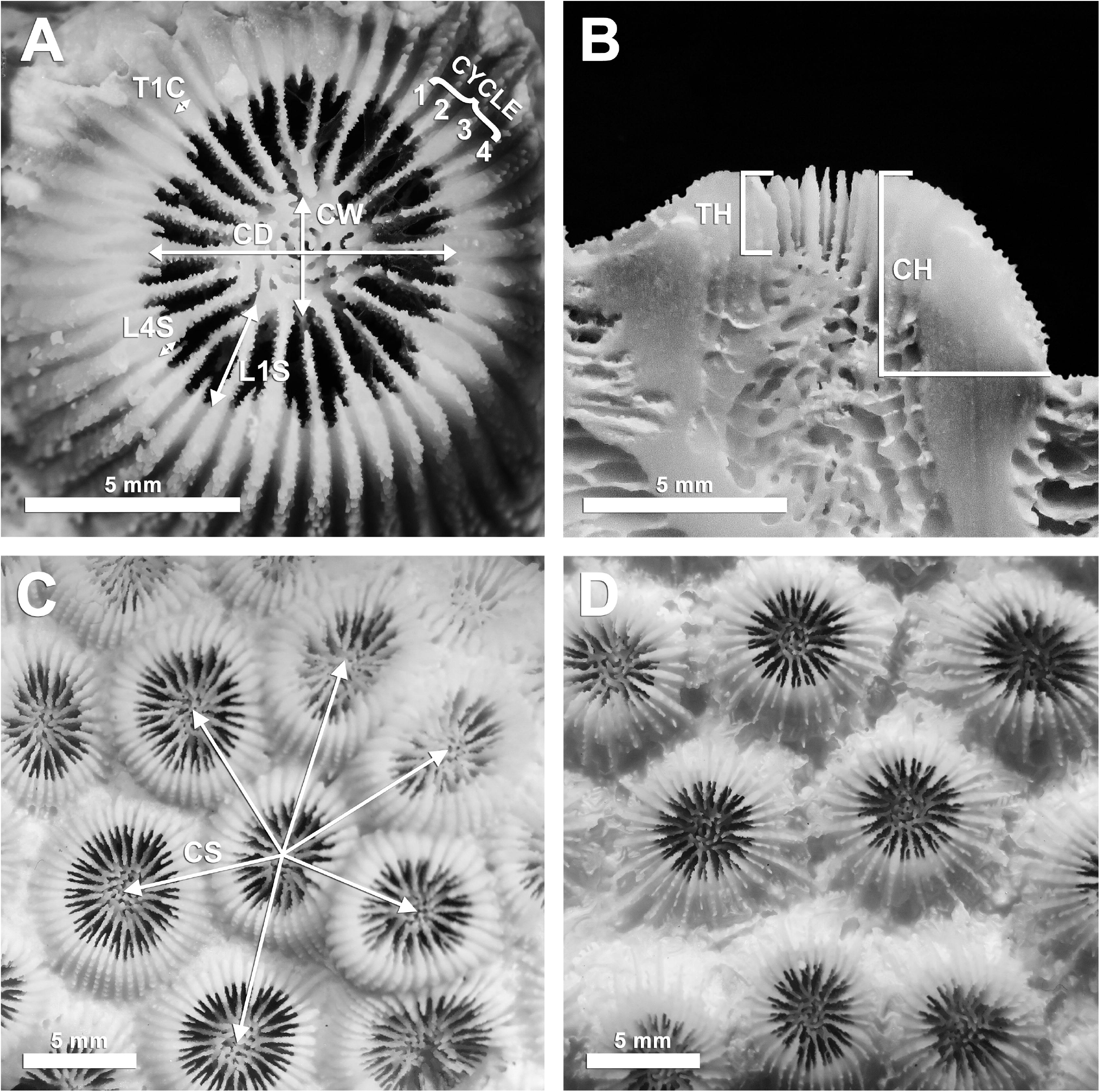
Corallite morphology photo panel. (A) Five of the eight morphometric characters chosen for this study superimposed over a *Montastraea cavernosa* corallite. (B) Vertical morphometric characters superimposed over a transverse section of a corallite. (C) Typical corallite appearance for ‘shallow’ morphotype, with corallite spacing character superimposed. (D) Typical corallite appearance for ‘depth-generalist’ morphotype. Panels (C) and (D) were photographed at equal scales. Character abbreviations as in Table 1.

**Table 1.**
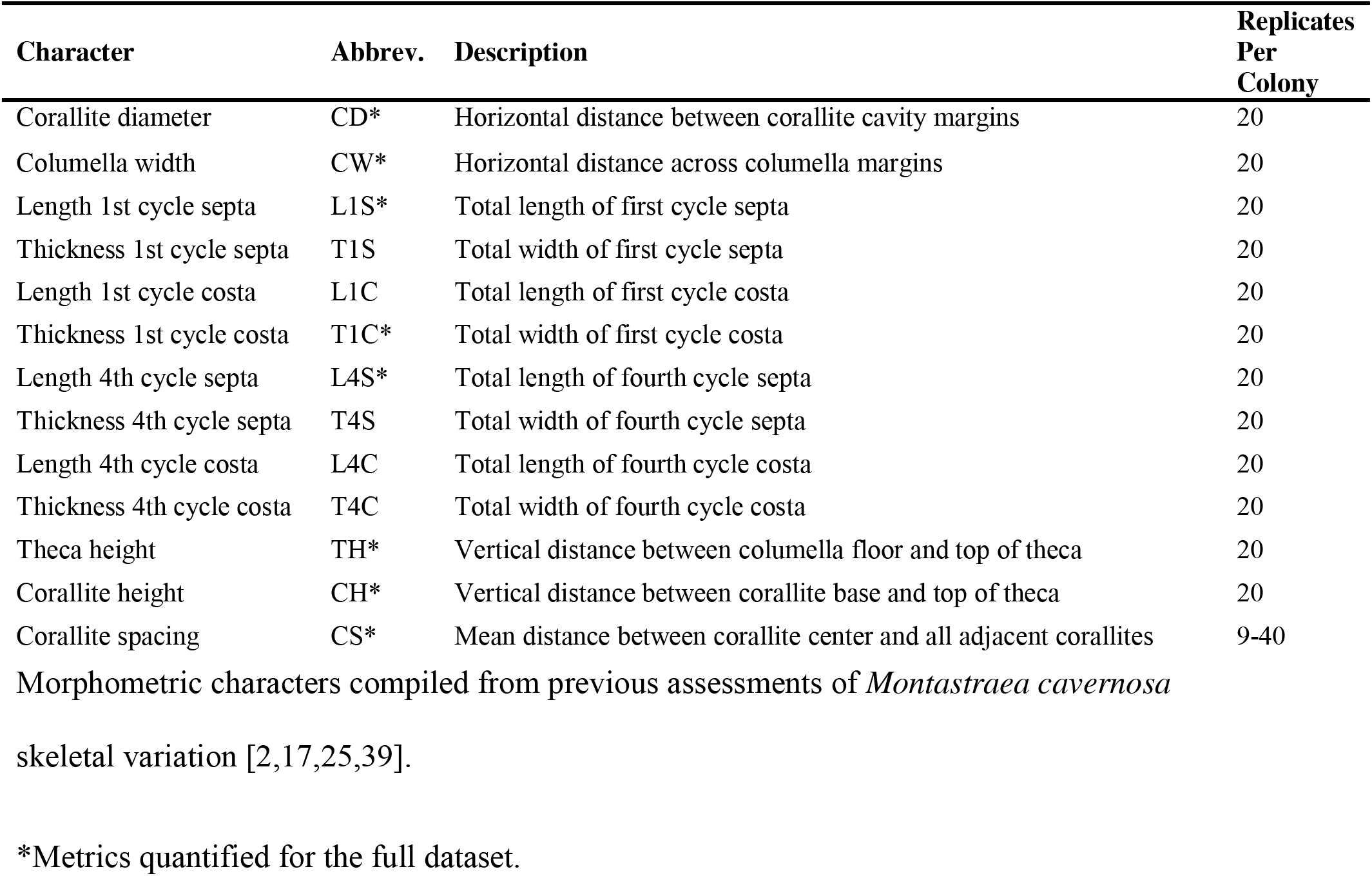
Corallite morphological characters.

### Statistical analyses

Means and standard deviations of each morphological character were calculated for coral samples and duplicate statistical analyses were conducted using each dataset to assess both intercolony and intracolony morphological variation. Coral sample data were analyzed using non-parametric tests due to violations of normality assumptions that could not be corrected via transformation. First, each morphological character was analyzed for significant variation across a single-factor combination of site and depth zone using Kruskal-Wallis tests. Pairwise comparisons were conducted using Dunn’s tests and *p* values were false discovery rate (FDR)- corrected in the R package *FSA* [42]. Due to the unbalanced sampling design, sample sets were tested for multivariate homogeneity of dispersions using the PERMDISP function in Primer v7 [43,44]. Assumptions of multivariate homogeneity of dispersions were violated (*p* < 0.05) for all datasets except for the standard deviation dataset, which can impact rejection rates for nonparametric statistical tests. However, the highest variance was associated with the largest sample sizes in all but the symbiont dataset, which likely increases the conservatism of the multivariate test results [45]. Two-way permutational multivariate analyses of variance (PERMANOVAs) tested the interactive effects of site (West FGB, East FGB, Bright, McGrail, Pulley Ridge, Dry Tortugas) and depth zone (mesophotic, shallow) on overall corallite morphology, including pairwise comparisons within factors. Additional pairwise comparisons were conducted for West and East FGB samples as an assessment of morphological variation between depth zones within sites. Test conditions for two-way PERMANOVAs utilized Euclidean distance, Type III SS, permutation of residuals under a reduced model, and 9999 model permutations in Primer v7 [43,44]. Subsequently similarity percentage (SIMPER) tests determined which morphometric characters contributed most strongly to differences observed among site and depth factor groups. SIMPER parameters utilized Euclidean distance and an 80% contribution cutoff.

Corallite morphological variation across all sites was visualized with principal coordinates analysis (PCoA) and two potential morphotypes were identified using sample groupings. Clustering patterns from the PCoA were tested for the presence of two morphotypes using a *k*-means clustering test (kRCLUSTER) with *k*=2. Frequency distributions were visualized for the six most variable characters across depth (corallite diameter, columella width, corallite spacing, theca height, corallite height, and length of the first cycle septa) to attempt to identify a threshold measurement that would allow a quantitative determination of morphotype from a single character. Morphotype assignments from the *k*-means clustering test were compared to assignments from the corallite spacing (CS) metric to determine assignment accuracy.

To assess which morphological characters differed between morphotypes while controlling for variation in environmental conditions across depth, a subset dataset was created using samples from the shallow caps (20 m) at West and East FGB. Additionally, corresponding algal symbiont (Symbiodiniaceae) density, areal chlorophylls *a* and *c2*, cellular chlorophylls *a* and *c2*, and chlorophyll *a*:*c2* ratio data from a recent study of the same coral samples [37,38] were added to the analyses. Symbiont and chlorophyll data were collected as described in Polinski and Voss [37]. This reduced dataset was tested for significant differences between ‘depth-generalist’ (n=20) and ‘shallow’ (n=46) morphotypes using Mann-Whitney U tests with the R package *ggpubr* [46]. Next, separate one-way PERMANOVA and SIMPER tests were conducted to determine if overall corallite morphology and overall symbiont/chlorophyll parameters were different between morphotypes sampled from the same depth zone.

Morphotype assignments for samples from West and East FGB were also matched to genotypes generated from the same colonies to identify any relationship between corallite morphology and genetic structure. Based on evidence of low genetic differentiation within the Flower Garden Banks [36], samples were combined from West and East FGB sites to form two populations based on morphotype assignments. Samples missing sufficient microsatellite marker coverage were removed from the analyses. Multi-locus genotypes were scored and normality assumptions were checked as described in Studivan and Voss [36] and assessment of population structure (with depth-generalist: n=70, shallow: n=35) was conducted with an analysis of molecular variance (AMOVA) using fixation index (F_ST_) in GenAlEx 6.5 [47,48] and with genetic structure analysis in Structure 2.3.4 and Structure Harvester [49–51].

## Results

### Corallite variation

Specimens from mesophotic zones (sites West FGB, East FGB, Bright, McGrail, and Pulley Ridge) had smaller mean corallite diameter (Kruskal-Wallis: *p*=3.5e^-8^), increased mean corallite spacing (*p=*2.0e^-16^), and larger mean corallite height (*p=*2.0e^-16^) compared to specimens from shallow zones (sites West FGB, East FGB, and Dry Tortugas; Table 2, Fig 2, S3 Table, S2 Fig). The two-way PERMANOVA of all samples revealed both site (Pseudo-F_5, 204_=14.99, *p=*0.0001) and depth zone (Pseudo-F_1, 204_=70.12, *p=*0.0001) as significant factors affecting corallite morphology across all metrics, while the interaction between site and depth zone was not significant (Pseudo-F_1, 204_=2.55, *p=*0.066). Pairwise PERMANOVA comparisons among sites within depth zones identified significant morphological variation primarily between both Pulley Ridge and Dry Tortugas compared to all other sites (Table S4). Pairwise comparisons between depths at West and East FGB showed a significant effect of depth zone on overall corallite morphology (West FGB: t=5.13, *p=*0.0001; East FGB: t=5.98, *p=*0.0001). Replicated statistical tests using standard deviation data corroborated the same trends (site: Pseudo-F_5, 204_=5.05, *p=*0.0001; depth: Pseudo-F_1, 204_=18.94, *p=*0.0001; interaction: Pseudo-F_1, 204_=0.29, *p=*0.89; S5 Table), indicating that intracolony variation was likely accounted for in the sampling and measurement design. SIMPER analyses revealed that corallite variation between shallow and mesophotic zones across all sites was primarily attributed to corallite spacing (73.15%) and corallite diameter (11.61%).

**Fig 2.**
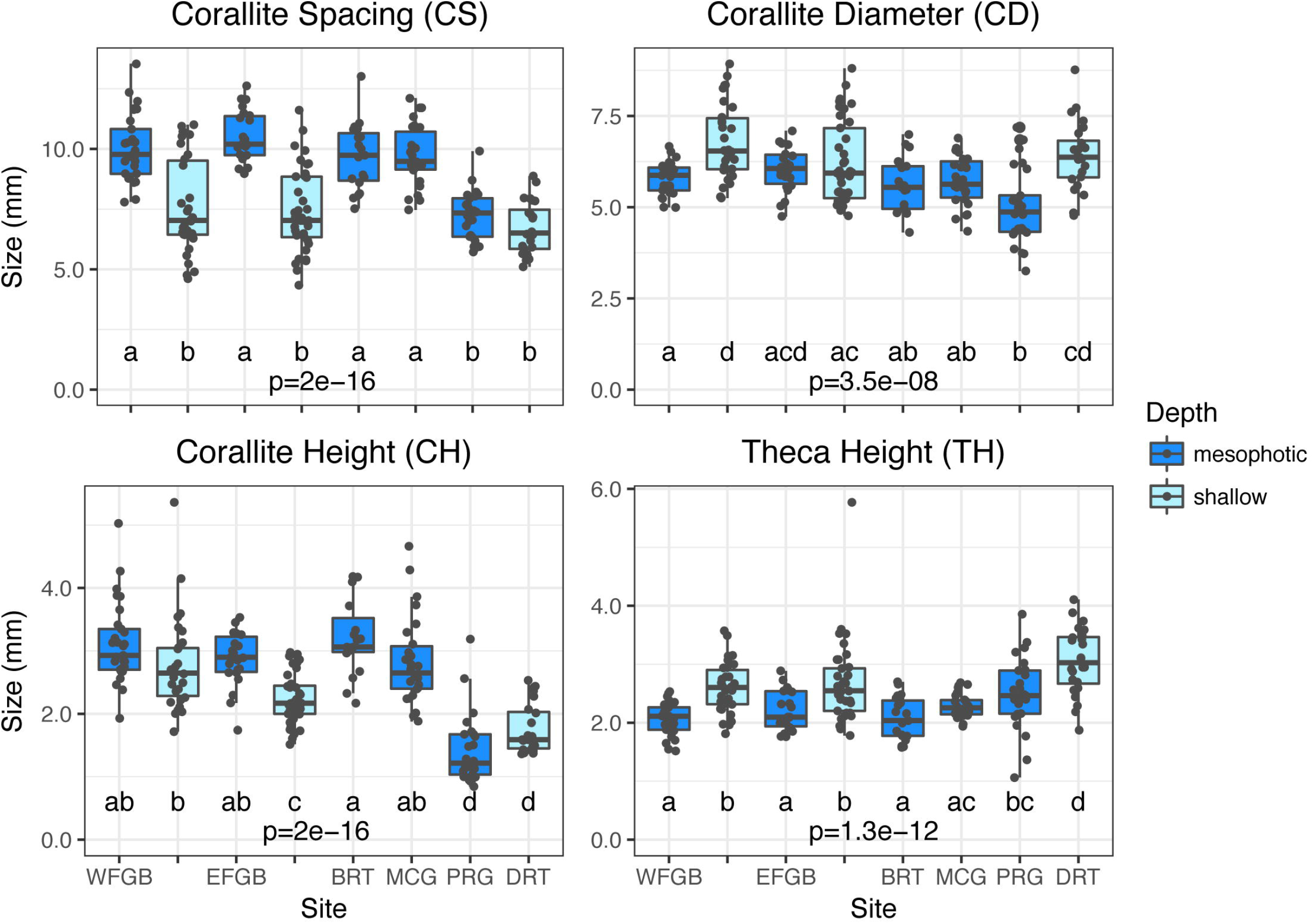
Corallite sample means. Boxplots with sample overlays for corallite spacing (CS), corallite diameter (CD), corallite height (CH), and theca height (TH) across six sites and two depth zones in the Gulf of Mexico. Overall *p* values represent Kruskal-Wallis tests across sites and depth zones for each metric and different letters denote significant differences (*p*<0.05) between pairwise comparisons of sites and depth zones generated by Dunn’s tests.

**Table 2.** Corallite sample means.

| Site | Depth Zone | Sampling Depth (m) | Sample Size (n) | Latitude* | Longitude* | CD | CW | L1S | L4S | T1C | TH | CH | CS |
| --- | --- | --- | --- | --- | --- | --- | --- | --- | --- | --- | --- | --- | --- |
| West FGB | mesophotic | 45 | 29 | 27.87510 | -93.82035 | 5.76 ± 0.09 | 2.32 ± 0.05 | 1.64 ± 0.03 | 0.51 ± 0.02 | 0.32 ± 0.01 | 2.06 ± 0.05 | 3.10 ± 0.11 | 10.06 ± 0.25 |
|  | shallow | 20 | 30 | 27.87495 | -93.81637 | 6.79 ± 0.18 | 2.88 ± 0.09 | 1.22 ± 0.07 | 0.39 ± 0.02 | 0.24 ± 0.01 | 2.60 ± 0.07 | 2.75 ± 0.13 | 7.59 ± 0.36 |
| East FGB | mesophotic | 45 | 22 | 27.91102 | -93.59668 | 6.02 ± 0.12 | 2.53 ± 0.06 | 1.81 ± 0.05 | 0.62 ± 0.03 | 0.34 ± 0.01 | 2.20 ± 0.07 | 2.87 ± 0.09 | 10.49 ± 0.22 |
|  | shallow | 20 | 37 | 27.90987 | -93.60021 | 6.22 ± 0.18 | 2.77 ± 0.08 | 1.18 ± 0.04 | 0.43 ± 0.01 | 0.28 ± 0.01 | 2.68 ± 0.11 | 2.24 ± 0.06 | 7.42 ± 0.28 |
| Bright Bank | mesophotic | 50 | 19 | 27.88620 | -93.30174 | 5.60 ± 0.17 | 2.38 ± 0.08 | 1.53 ± 0.06 | 0.49 ± 0.02 | 0.33 ± 0.01 | 2.07 ± 0.08 | 3.19 ± 0.13 | 9.68 ± 0.31 |
| McGrail Bank | mesophotic | 50 | 26 | 27.96299 | -92.59262 | 5.70 ± 0.12 | 2.27 ± 0.04 | 1.66 ± 0.05 | 0.51 ± 0.02 | 0.32 ± 0.01 | 2.28 ± 0.03 | 2.82 ± 0.13 | 9.71 ± 0.24 |
| Pulley Ridge | mesophotic | 65 | 25 | 24.79382 | -83.67401 | 4.99 ± 0.20 | 2.24 ± 0.08 | 0.84 ± 0.02 | 0.29 ± 0.01 | 0.18 ± 0.01 | 2.50 ± 0.12 | 1.41 ± 0.11 | 7.23 ± 0.20 |
| Dry Tortugas | shallow | 30 | 24 | 24.47279 | -82.96807 | 6.41 ± 0.18 | 3.04 ± 0.06 | 0.94 ± 0.03 | 0.35 ± 0.01 | 0.23 ± 0.01 | 3.05 ± 0.11 | 1.75 ± 0.07 | 6.73 ± 0.22 |

Mean ± standard error of the mean (SEM) in millimeters of eight corallite morphometrics across six sites and two depth zones in the Gulf of Mexico.
\*Geographic coordinates given as decimal degrees (WGS84).

Principal coordinates analysis clustered samples into two groups (Fig 3). One group consisted of the majority of samples from Bright and McGrail Banks, while a second group consisted primarily of Dry Tortugas samples. The split between groups did not lie between shallow and mesophotic samples, however. Corals from West and East FGB were found in both sample groupings, identifying a dichotomy in overall corallite morphology (i.e. two morphotypes). A subset of samples collected at 20 m from both FGB sites had overall corallite structure more similar to mesophotic samples than to the remaining shallow samples. Pulley Ridge samples appeared to share some morphological characteristics of both morphotypes, indicated by overlap between both sample clusters in the PCoA (Fig 3). Despite having smaller mean corallite diameters typical of other mesophotic corals (Fig 2), the Pulley Ridge colonies demonstrated corallite spacing consistent with the morphotype observed most commonly in shallow zones of both FGB sites and Dry Tortugas. With the exception of Pulley Ridge, the majority of mesophotic corals appeared to be a single morphotype (mesophotic West and East FGB, Bright, and McGrail), while shallow corals were split between both morphotypes.

**Fig 3.**
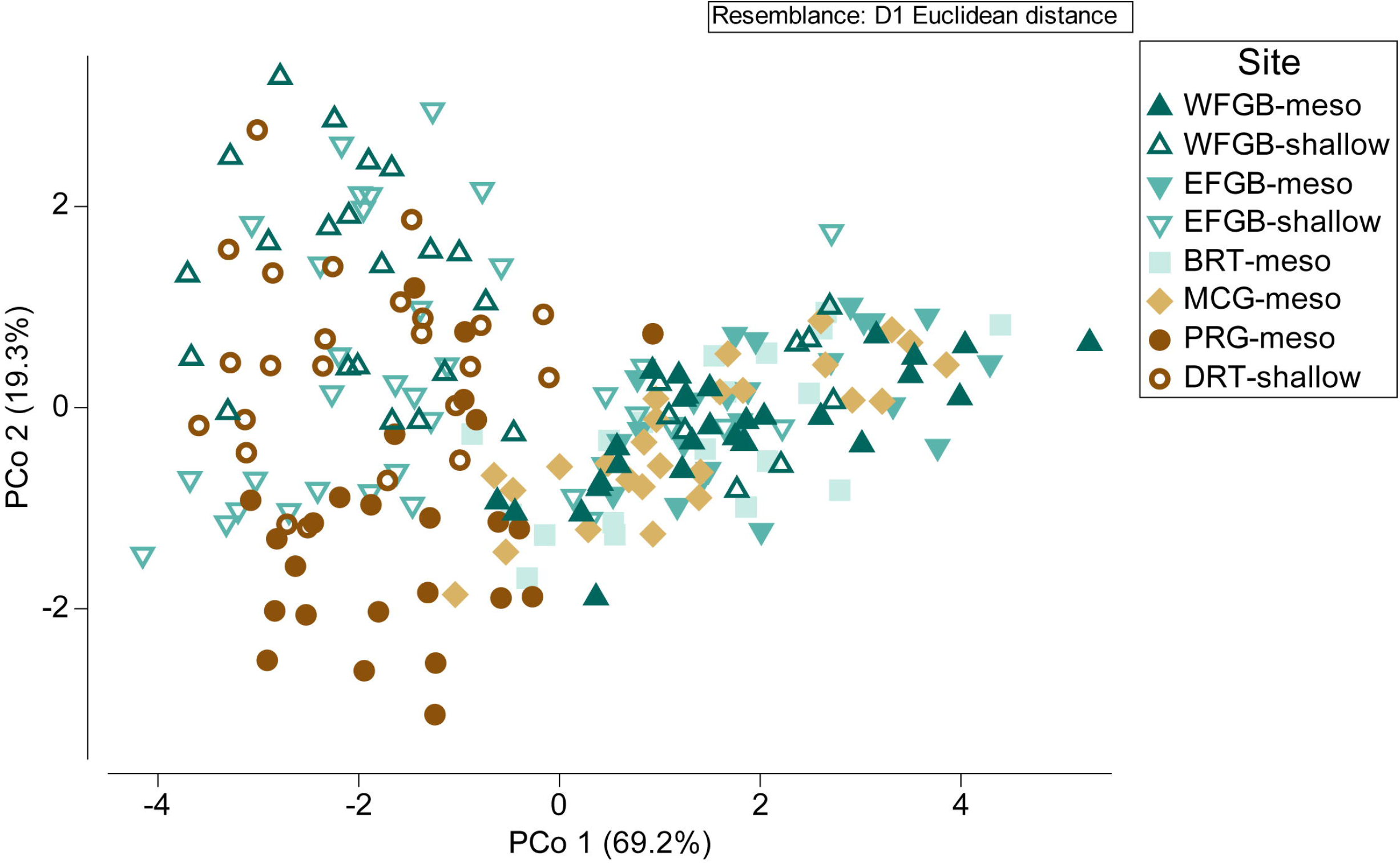
Principal coordinates analysis. Principal coordinates analysis (PCoA) ordination of corallite morphology across six sites in the Gulf of Mexico, explaining 88.5% of the total variation (PCo 1: 69.2%, PCo 2: 19.3%). Degree of difference among samples represented by Euclidean distance between sample points. Color and shape of each point corresponds to site and depth zone.

### Morphotype assignment

Frequency distributions of mean corallite diameter (CD), width (CW), spacing (CS), height (CH), theca height (TH), and first cycle septal length (L1S) revealed relatively unimodal distributions for most metrics except for corallite spacing and first cycle septal length (S3 Fig). The bimodal distribution observed in mean corallite spacing across all samples was indicative of two morphotypes split between 8.2–8.6 mm. All samples were sorted by corallite spacing and assigned a morphotype designation where CS>8.4 mm was considered ‘depth-generalist’ and CS<8.4 mm was considered ‘shallow.’ Morphotype assignments using the corallite spacing method were found to have a correct assignment rate of 93.87% when compared to assignments from the *k*-means clustering method (Fig 4; kRCLUSTER: R=0.746, 13 incorrect out of 212 assignments). Following the patterns observed in the PCoA (Fig 3), the majority of samples collected from mesophotic depth zones at West and East FGB, Bright, and McGrail were identified as the depth-generalist morphotype, while most samples from Dry Tortugas and Pulley Ridge were identified as the shallow morphotype. Samples collected from the shallow depth zone at West and East FGB included both morphotype assignments.

**Fig 4.**
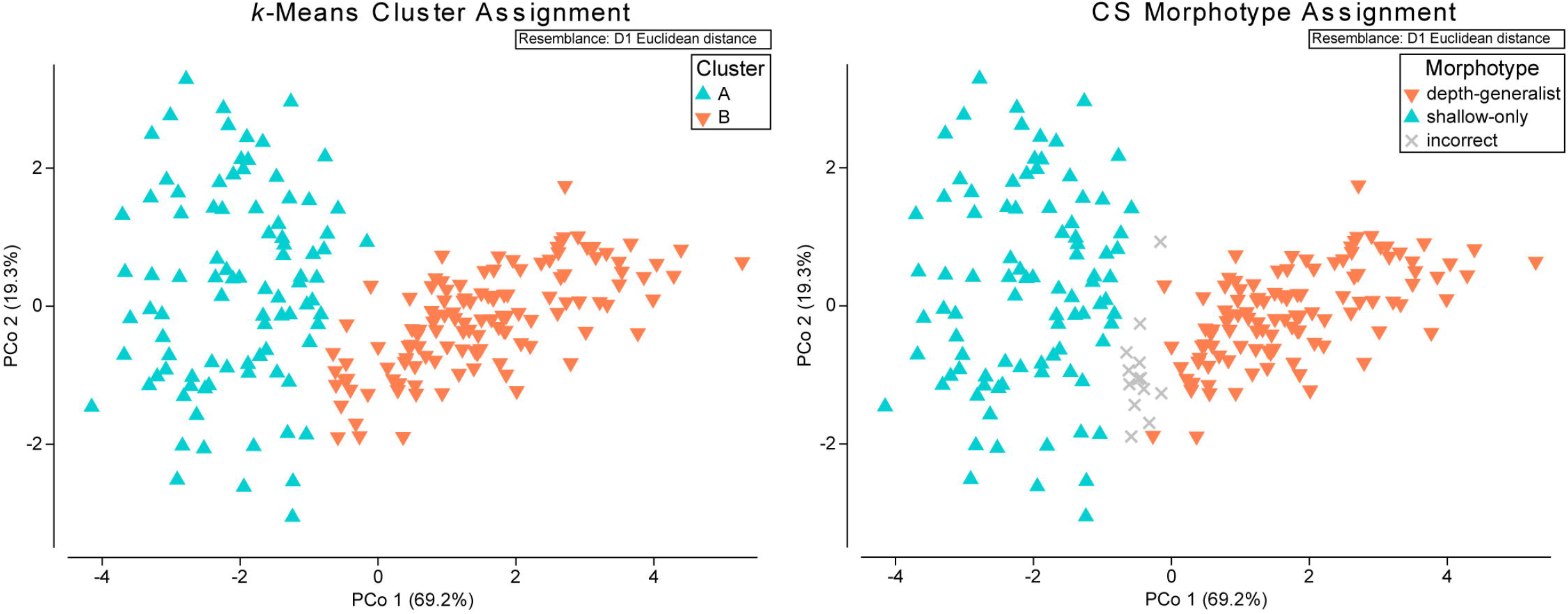
Comparison of morphotype assignment methods. Principal coordinates analyses (PCoA) of corallite morphology across six sites in the Gulf of Mexico, with color overlays corresponding to morphotypes. Morphotype assignments were made using a *k*-means cluster test (left; kRCLUSTER: R=0.746) and by the threshold in corallite spacing measurements at 8.40 mm (right). Assignments using the corallite spacing method had a correct assignment rate of 93.87% when compared to assignments from the *k*-means clustering method (13 incorrect out of 212 assignments, shown in gray).

Morphotype assignments from the *k*-means clustering method were used to create subset datasets of samples collected in the shallow depth zones (20 m) of West and East FGB and tested whether the two morphotypes had significantly different corallite structure in the same environment. The differences among morphotypes were not solely attributed to morphological variation due to depth. The depth-generalist morphotype found at 20 m demonstrated increased mean corallite spacing (Mann-Whitney U: *p*=1.6e^-10^) and reduced corallite diameters (*p*=0.0056), meaning there were fewer and smaller corallites per unit area as compared to the shallow morphotype. The depth-generalist morphotype also had taller corallites (corallite height: *p*=9.1e^-5^) and notably longer first septae (length of first cycle septa: *p*=5.6e^-10^) (Table 3, Fig 5, S4 Fig). A single factor PERMANOVA using morphotype assignment identified a significant difference between depth-generalist and shallow morphotypes (Pseudo-F_1, 65_=62.73, *p=*0.0001). For the SIMPER analysis of morphotypes, corallite spacing (70.93%) and diameter (13.74%) were the most influential contributors to variation between morphotypes. Mann-Whitney U tests indicated that mean symbiont density (Mann-Whitney U: *p=*0.0028), areal chlorophylls *a* (*p*=1.7e^-8^) and *c2* (*p*=7.2e^-8^), and cellular chlorophylls *a* (*p*=5.4e^-5^) and *c2* (*p*=0.0051) were significantly higher in the depth-generalist morphotype compared to the shallow morphotype (Table 3, S5 Fig), despite being found in a similar light regime at 20 m and with nearly-identical Symbiodiniaceae community assemblages [37]. The single factor PERMANOVA identified a multivariate difference between depth-generalist and shallow morphotypes across symbiont and chlorophyll metrics (Pseudo-F_1, 65_=11.329, *p=*0.0016), and the SIMPER attributed 100% of the variation between morphotypes to symbiont density.

**Fig 5.**
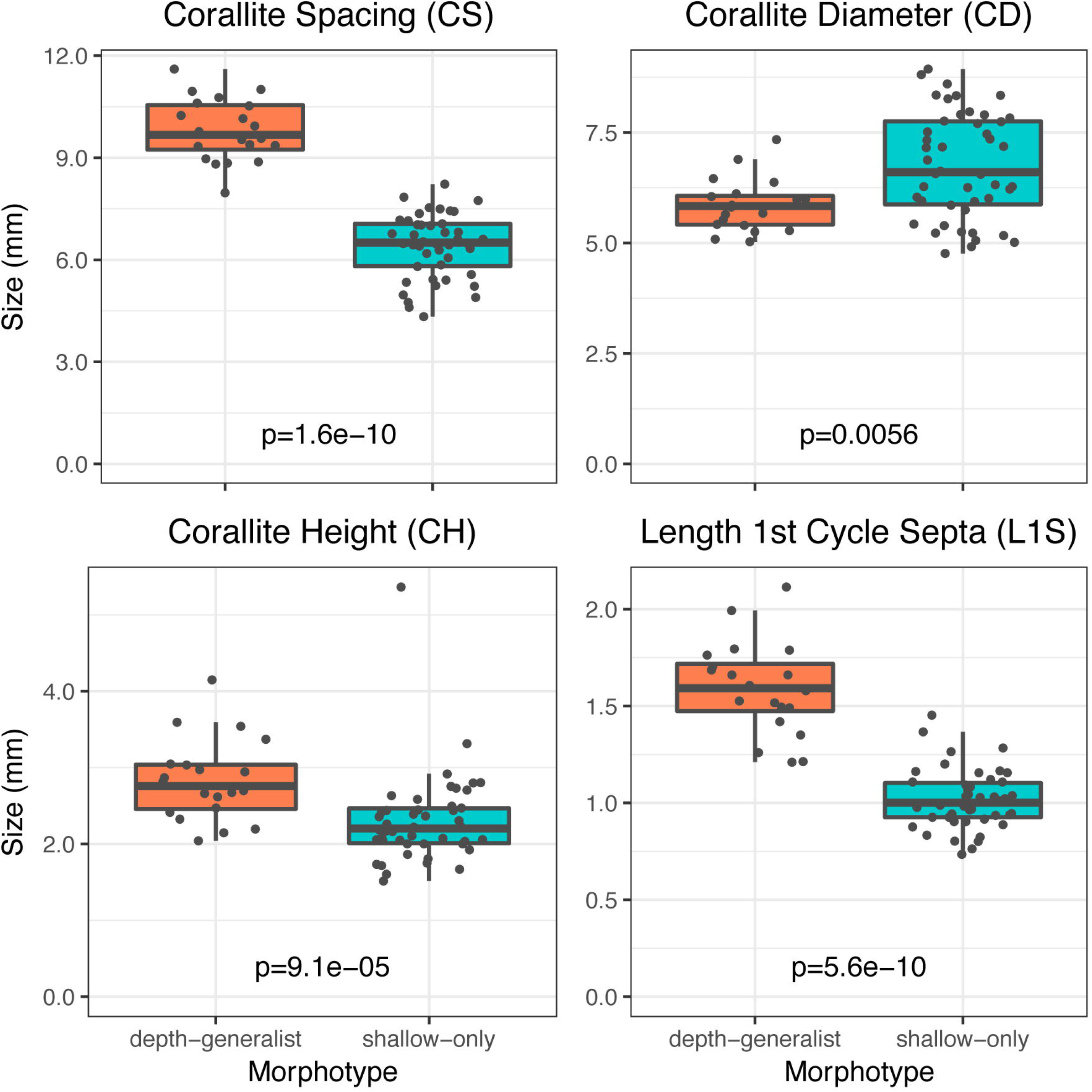
Morphotype sample means. Boxplots with sample overlays for corallite spacing (CS), corallite diameter (CD), corallite height (CH), and length of first cycle septa (L1S) across depth-generalist (n=20) and shallow (n=46) morphotypes sampled within the shallow zone of West and East FGB. Overall *p* values represent Mann-Whitney U tests between morphotype for each metric.

**Table 3.** Morphotype sample means.

| Morphotype | Sample Size (n) | CD | CW | L1S | L4S | T1C | TH | CH | CS | Areal Chl (ug cm<sup>-2</sup> ± SEM) |  | Cellular Chl (pg cell<sup>-1</sup> ± SEM) |  | Chl *a:c<sub>2</sub>* | Zooxanthellae Density (millions cells cm<sup>-2</sup> ± SEM) |
| --- | --- | --- | --- | --- | --- | --- | --- | --- | --- | --- | --- | --- | --- | --- | --- |
|  |  |  |  |  |  |  |  |  |  | chl *a* | chl *c<sub>2</sub>* | chl *a* | chl *c<sub>2</sub>* |  |  |
| depth-generalist | 20 | 5.85 ± 0.13 | 2.48 ± 0.06 | 1.59 ± 0.05 | 0.49 ± 0.02 | 0.34 ± 0.01 | 2.74 ± 0.18 | 2.82 ± 0.11 | 9.81 ± 0.20 | 11.13 ± 0.62 | 4.35 ± 0.44 | 7.05 ± 0.62 | 2.75 ± 0.31 | 2.71 ± 0.10 | 1.73 ± 0.12 |
| shallow | 46 | 6.75 ± 0.17 | 2.97 ± 0.07 | 1.01 ± 0.02 | 0.38 ± 0.01 | 0.23 ± 0.01 | 2.60 ± 0.06 | 2.30 ± 0.08 | 6.42 ± 0.13 | 6.42 ± 0.20 | 2.45 ± 0.08 | 4.86 ± 0.16 | 1.85 ± 0.06 | 2.64 ± 0.04 | 1.36 ± 0.04 |

Mean ± standard error of the mean (SEM) of eight measured corallite morphometrics, algal symbiont density, and chlorophyll concentrations across depth-generalist and shallow morphotypes from within the shallow zone of West and East FGB.

Within the samples from West and East FGB, comparison of population structure using morphotype as a factor revealed that corallite variation had a weak relationship to genetic variation. The fixation index (F_ST_), which represents the level of genetic differentiation between populations, was low but demonstrated significant genetic structure between depth-generalist and shallow morphotypes at both FGB sites combined (AMOVA: F_ST_=0.007, *p*=0.008). A second AMOVA using morphotype assignments according to the corallite spacing method also demonstrated significant genetic structure (AMOVA: F_ST_=0.007, *p*=0.005). However, structure analyses with both assignment methods indicated that there was one genetic cluster (K=1, panmixia) found at both FGB sites (S6 Table, S6 Fig), which is reflective of the results of a larger genotypic examination of coral populations in the NW GOM [36].

## Discussion

### Variation among shallow and mesophotic *M. cavernosa*

Previous studies have observed that mean corallite diameter decreases and mean corallite spacing increases with diminishing light in some scleractinian species [2,6,9,20,25,52,53]. This study extends this observation to mesophotic zones with *M. cavernosa* (see also [69]), as mesophotic corallites were on average smaller and further apart than their shallow counterparts. Mesophotic corallites were also typically taller (corallite height), but internally shallower (theca height) than shallow corallites (Fig 2). Morphological variation across depth zones may be the result of corals’ physiological responses to environmental conditions in mesophotic zones, including maximizing tissue area for light capture while minimizing self-shading and metabolic costs [11,23,53,54], and/or maximizing tentacle area exposed for food capture [55]. While colonies examined in this study were not quantitatively observed for polyp extension or feeding behavior, earlier studies with *M. cavernosa* hypothesized that smaller corallites may also correspond with increased polyp opening, although the tradeoffs between photosynthetic and heterotrophic yields are not well understood [11,56]. Increased algal symbiont density and chlorophyll concentration in these same colonies is consistent with mesophotic corals demonstrating unique photoadaptive strategies in low light environments compared to shallow conspecifics [37,38]. Mesophotic *M. cavernosa* from the Flower Garden Banks and Pulley Ridge contained more symbionts and chlorophylls *a* and *c2* than their shallow conspecifics from the Flower Garden Banks and Dry Tortugas, respectively.

While increased pigmentation and symbiont densities are common photoadaptive responses to lower-light environments [52,57–59], they are suspected to also reduce light penetration to deeper coral tissues due to greater optical thickness via self-shading [52]. Furthermore, a recent study identified photoconvertible red fluorescent proteins that transform poorly-absorbed blue-green light to orange-red wavelengths for increased light absorption, which were found more commonly in mesophotic corals [60]. However, while these strategies may maximize light absorption, they likely result in lower tissue penetration. Corals in light-limited environments, such as mesophotic coral ecosystems, may mitigate the negative effects of photoadaptation with enhanced light scattering from skeletal structures [58,61]. Flat surfaces, such as flattened mesophotic colony skeletons, can increase light scattering threefold within tissues, with additional enhancement caused by concave surfaces in the interior of corallites and by complex structures such as septa [62–65]. This study identified an increase in mean septal length (see also [2]) and corallite height of mesophotic samples (Fig 2), which may provide increased light scattering at depth. It is likely that a tradeoff exists between shallow and mesophotic morphologies, given multiple responses at the symbiotic and skeletal levels, although their effect on coral physiology and intraspecies competition is not well known at this time.

### Identification of ‘depth-generalist’ and ‘shallow’ morphotypes

The *M. cavernosa* morphotypes described here appear to differ somewhat from the nocturnal and diurnal morphotypes identified in previous studies [11–14,17]. Previous examination of *M. cavernosa* corallite structure has characterized morphotypes by variation in corallite size and tentacle behavior (open versus closed polyps). The study presented here identified corallite spacing as the primary determinant of morphotype, with corallite size as significant, but less important, factor driving multivariate differences between the two morphotypes. Mean corallite diameters of both morphotypes were comparable to previous studies (Table 3), but we did not observe obvious differences in tentacle behavior between morphotypes. Rather, it was common to find colonies with a portion of the polyps open while the others remained closed. In one previous study, nocturnal and diurnal morphotypes were distributed differently over depth, with the smaller-corallite diurnal morphotype more common in habitats <10 m and the larger nocturnal morphotype found primarily at 15–25 m [17]. A similar study identified two comparable morphotypes across the TWA with distinct variation in corallite diameter, but found no apparent depth distribution between 5–22 m in Belize [15]. The present study identified a smaller-corallite morphotype with a broader depth distribution (20–65 m), while a larger-corallite morphotype was mostly restricted to shallower habitats (20–30 m; except at Pulley Ridge, see below). The discrepancies regarding morphotype identification among these studies suggest that beyond feeding strategies or depth alone, additional genotypic or external factors are likely impacting the observed *M. cavernosa* morphologies. Whereas Ruiz Torres [17] suspected that differences between morphotypes may have constituted cryptic speciation within *M. cavernosa*, we observed limited genetic differentiation between morphotypes (see also [15]).

Based on the observed depth distribution of morphological variants across six sites examined in the GOM, we identified distinct ‘depth-generalist’ and ‘shallow’ morphotypes within *M. cavernosa* (Fig 4). Differences in overall corallite morphologies were mainly attributed to increased corallite spacing in the depth-generalist type (Fig 1C and 1D), but there were also significant differences in corallite diameter, corallite and theca height, and septal length. The depth-generalist morphotype was characterized by smaller and more widely-spaced corallites that were taller over the surrounding skeleton (Fig 5). We observed that the variation in corallite spacing alone was enough to predict the correct morphotype assignment with a relatively high level of accuracy (93.87%) compared to a cluster analysis of morphological variation across all eight characters (Fig 4). This is of potential interest for rapid, non-destructive identification of *M. cavernosa* morphotypes *in situ*, perhaps allowing targeted sampling of morphotypes for comparative genotyping analyses across broad spatial scales. Since assignment accuracy using the corallite spacing method was also consistently high among distant sites in the GOM (∼1,000 km separation), the potential for widespread presence of these morphotypes should be explored further throughout this species’ range.

### Potential genotypic influence on morphotype

The comparison of morphotypes with genetic structure within the Flower Garden Banks determined that morphotype, regardless of assignment method, had a small but significant effect on population differentiation. Despite evidence that morphotypes were significantly differentiated, structure analysis predicted population panmixia (see also [36]). It appears that the slight differentiation is the result of genotypic differences between morphotypes, although there was no evidence to suggest cryptic speciation or a lack of gene flow between morphotypes. However, it must be noted that analyses using nine microsatellite loci (this study) or nuclear markers *ß-tubulin* and mitochondrial marker *cox1* [15,66] may not be as sensitive to detecting cryptic morphotypes or identifying selection as analyses using many markers including amplified fragment length polymorphism (AFLP) loci [67] or single nucleotide polymorphism (SNP) loci [68]. Results from previous population genetics studies suggest that there is a high degree of polymorphism within *M. cavernosa* populations across the Tropical Western Atlantic through ecological timescales [15,16], yet without strong evidence of genetic isolation among morphotypes. Polymorphic differences may represent physiological tradeoffs among individuals pertaining to differences in feeding strategy, calcification, optical properties and light capture, symbiont communities, and perhaps even stress resilience [9,11,58,65]. The depth-generalist morphotype demonstrated skeletal characteristics that may contribute to enhanced light capture in mesophotic zones through increased light scatter within the skeleton, including taller corallites and longer septa. The depth-generalist morphotype also had higher mean symbiont densities and chlorophyll concentrations (S5 Fig) that likely support increased light capture [11]. Corallite characteristics and increased symbiont densities were retained in the depth-generalist morphotype even in shallow zones, suggesting aspects of the symbiosis other than light availability control the abundance of algal cells and chlorophyll content.

During high light conditions leading to thermal stress, increased light amplification can result in more severe bleaching responses [69]. As symbiont density is reduced through bleaching, the effect of light scattering increases and creates a positive feedback loop, exposing additional symbionts to high intensities of light [65]. As a result, corals with higher light scattering skeletal properties may confer lower bleaching resistance and be therefore less suited to higher light environments. Bleaching events due to excess solar radiation may be more likely in shallower reef habitats as compared to mesophotic habitats [28]. While the relatively high-latitude Flower Garden Banks are not typically exposed to high temperature stressors seen elsewhere in the Tropical Western Atlantic, four bleaching events have been observed in the last three decades [70]. We hypothesize that the depth-generalist morphotype may therefore be less abundant in shallow zones of the Flower Garden Banks due in part to lower bleaching resistance.

The exclusion of the shallow morphotype from mesophotic sites was consistent across sites in the NW GOM (Fig 3), however, this pattern did not hold in the SE GOM, as most mesophotic *M. cavernosa* at Pulley Ridge were identified as the shallow morphotype and the depth-generalist morphotype was conspicuously absent from Dry Tortugas. To potentially explain the distribution of morphotypes in the SE GOM, we must also consider genetic variation beyond that attributed to morphological differences. Given the patterns of morphotype depth distribution in the NW GOM, Pulley Ridge would be expected to be primarily comprised of the depth-generalist morphotype and Dry Tortugas would be expected to include a mixed population of both morphotypes. The relative lack of the depth-generalist morphotype observed at both sites may have instead resulted from low larval dispersal and population connectivity in the SE GOM. Population genetics analyses suggest relative isolation of Pulley Ridge *M. cavernosa* from other sites in the GOM and Tropical Western Atlantic [32,36]. It is possible that Pulley Ridge and Dry Tortugas were initially colonized by the shallow morphotype, or alternatively, both morphotypes may have been recruited but environmental conditions in the region may favor corals with corallite spacing similar to the shallow morphotype. The former seems more likely given the relative isolation of the SE GOM and the small population size of *M. cavernosa* at Pulley Ridge estimated from surveys and migration models [36,71,72]. Divergence from the shallow morphotype at Pulley Ridge may have then been the result of photoadaptation towards smaller corallites typically found in mesophotic zones. Pulley Ridge corals exhibited significant deviation in corallite structure (notably corallite diameter, columella width, and theca height) from the shallow morphotype (Fig 2). Variations of corallite size and height were indicative of differences seen across shallow and mesophotic zones elsewhere in the GOM and likely represent skeletal adaptation among the mesophotic corals at Pulley Ridge to corallite characteristics better suited to low-light environments.

Throughout the fossil record of the past 25 million years, *M. cavernosa* populations in other regions of the Tropical Western Atlantic have demonstrated remarkable and persistent morphological variation that has not been attributed to any distinct genetic lineages [15,16]. The ability of multiple morphotypes to be maintained in absence of selection, reproductive isolation, or cryptic speciation may be due in part to high genetic diversity across much of this species’ range [73]. The results from this study reinforce the notion that morphological variation among *M. cavernosa* represents a combination of genotypic variation and phenotypic plasticity rather than responses to environmental stimuli alone. Additional assessments of skeletal optical properties and measures of photosynthetic performance could help determine whether morphotypes in *M. cavernosa* confer physiological and/or resilience tradeoffs. Characterizing such trade-offs may further elucidate the mechanisms that allow depth-generalist coral species to be successful across a variety of reef habitats.

## Acknowledgements

We acknowledge the staff of Flower Garden Banks and Florida Keys National Marine Sanctuaries; crews of the R/V *Manta* and M/V *Spree*; and L. Horn and J. White from the University of North Carolina at Wilmington Undersea Vehicle Program for assistance with sample collection. J. Beal, J. Emmert, J. Polinski, D. Dodge, A. Alker, P. Gardner, R. Susen, M. Ajemian, M. McCallister, C. Ledford, R. Christian, and M. Dickson provided diving support. Special thanks to J. Polinski, D. Perez, H. Clinton, P. Gardner, and D. Dodge for providing the algal symbiont and chlorophyll data, and to A. Alker for assisting with the morphometrics data collection. This is contribution number 2197 from Harbor Branch Oceanographic Institute at Florida Atlantic University.

## Supporting information

**S1 Fig. Map of the Gulf of Mexico with sampling sites.** Map of the Gulf of Mexico, with inset boxes of six sampling sites in the northwest and southeast Gulf of Mexico (NW GOM and SE GOM, respectively). Inset overlays include available bathymetry data of sites, and locations of specimen collection color-coded by depth zone (mesophotic 30-70 m, shallow 20-30 m). Geographic coordinates as in Table 2.

**S1 Table. Preliminary morphological character correlation matrix.** Correlation matrix of original thirteen corallite metrics compared across five preliminary coral samples collected from Dry Tortugas over a depth range of 25-33 m.

* Corallite spacing (CS) not examined. Insignificant correlations are shown as ns.

**S2 Table. Preliminary PERMANOVA results.** Test results for permutational multivariate analysis of variance (PERMANOVA, Overall) and univariate analyses (Kruskal-Wallis) across five preliminary coral samples collected from Dry Tortugas over a depth range of 25-33 m.

*Corallite spacing (CS) not examined. Insignificant *p* values are shown as ns.

**S3 Table. Kruskal Wallis results.** Test results for overall (Kruskal-Wallis) and pairwise comparisons (Dunn’s) across sites and depth zones for each morphological character.

*Insignificant pairwise comparisons denoted as ns.

**S2 Fig. Corallite sample means.** Boxplots with sample overlays for columella width (CW), thickness of the first cycle costa (T1C), length of the first cycle septa (L1S), and length of the fourth cycle septa (L4S) across six sites and two depth zones in the Gulf of Mexico. Overall *p* values represent Kruskal-Wallis tests across sites and depth zones for each metric and different letters denote significant differences (*p*<0.05) between pairwise comparisons of sites and depth zones generated by Dunn’s tests.

**S4 Table. PERMANOVA results.** Test results for permutational multivariate analysis of variance (PERMANOVA, Overall) and pairwise comparisons across sites within depth zones and across depth zones within sites.

*Insignificant pairwise comparisons denoted as ns.

**S5 Table. Standard deviation PERMANOVA results.** Test results for permutational multivariate analysis of variance (PERMANOVA) of intra-colony standard deviation and pairwise comparisons across sites within depth zones, and across depth zones within sites.

*Insignificant pairwise comparisons denoted as ns.

**S3 Fig. Size distributions of dominant characters.** Frequency distributions for the six morphological metrics representing the majority of corallite variation across depth, including: corallite diameter (CD), columella width (CW), corallite spacing (CS), theca height (TH), corallite height (CH), and length of first cycle septa (L1S). The size threshold of CS (8.40 mm) is represented in the orange vertical line, denoting two morphotypes distinguished primarily by differences in corallite spacing. L1S also had a split distribution but had a less obvious threshold (1.15 mm).

**S4 Fig. Morphotype sample means.** Boxplots with sample overlays for columella width (CW), thickness of the first cycle costa (T1C), theca height (TH), and length of fourth cycle septa (L4S) across depth-generalist (n=20) and shallow (n=46) morphotypes sampled within the shallow zone of West and East FGB. Overall *p* values represent Mann-Whitney U tests between morphotype for each metric.

**S5 Fig. Morphotype symbiont and chlorophyll means.** Boxplots with sample overlays for symbiont density, chlorophyll *a:c2*, areal chlorophyll *a*, areal chlorophyll *c2*, cellular chlorophyll *a*, and cellular chlorophyll *c2* across depth-generalist (n=20) and shallow (n=46) morphotypes. Overall *p* values represent Mann-Whitney U tests between morphotype for each metric.

**S6 Table. Evanno method for genetic structure.** Table describing the process behind population cluster (K) selection in structure analysis of depth-generalist and shallow populations using the *k*-means clustering method. Ten replicate structure models were run across a range of K values from 1–5 and model log likelihoods were compared. The Evanno method was used to determine the most likely number of K by identifying the largest change in likelihood (|Ln’’(K)|) and by comparing model probabilities in conjunction with variance (Delta K).

*Delta K not calculated for K=1 or K=5, so model likelihood was solely used to determine the most likely number of genetic clusters. The most likely number of K shown in bold.

**S6 Fig. Evanno method for genetic structure.** Plots describing the process behind population cluster (K) selection in structure analysis of depth-generalist and shallow populations using the *k*-means clustering method. Ten replicate structure models were run across a range of K values from 1–5 and model log likelihoods were compared. The Evanno method was used to determine the most likely number of K by identifying the largest change in likelihood (L(K)) and by comparing model probabilities in conjunction with variance (Delta K). Error bars represent standard deviation of the mean.

**S1 Dataset. Spreadsheet containing raw data.** Individual sheets for preliminary samples, raw corallite measurements, colony means, colony standard deviation, morphotype identification, symbiont metrics, genotypes for *k*-means morphotypes, and genotypes for CS morphotypes.

