## Supplementary figures and images for "*Montastraea cavernosa* corallite structure demonstrates distinct morphotypes across shallow and mesophotic depth zones in the Gulf of Mexico"

### Supplemental Fig 1

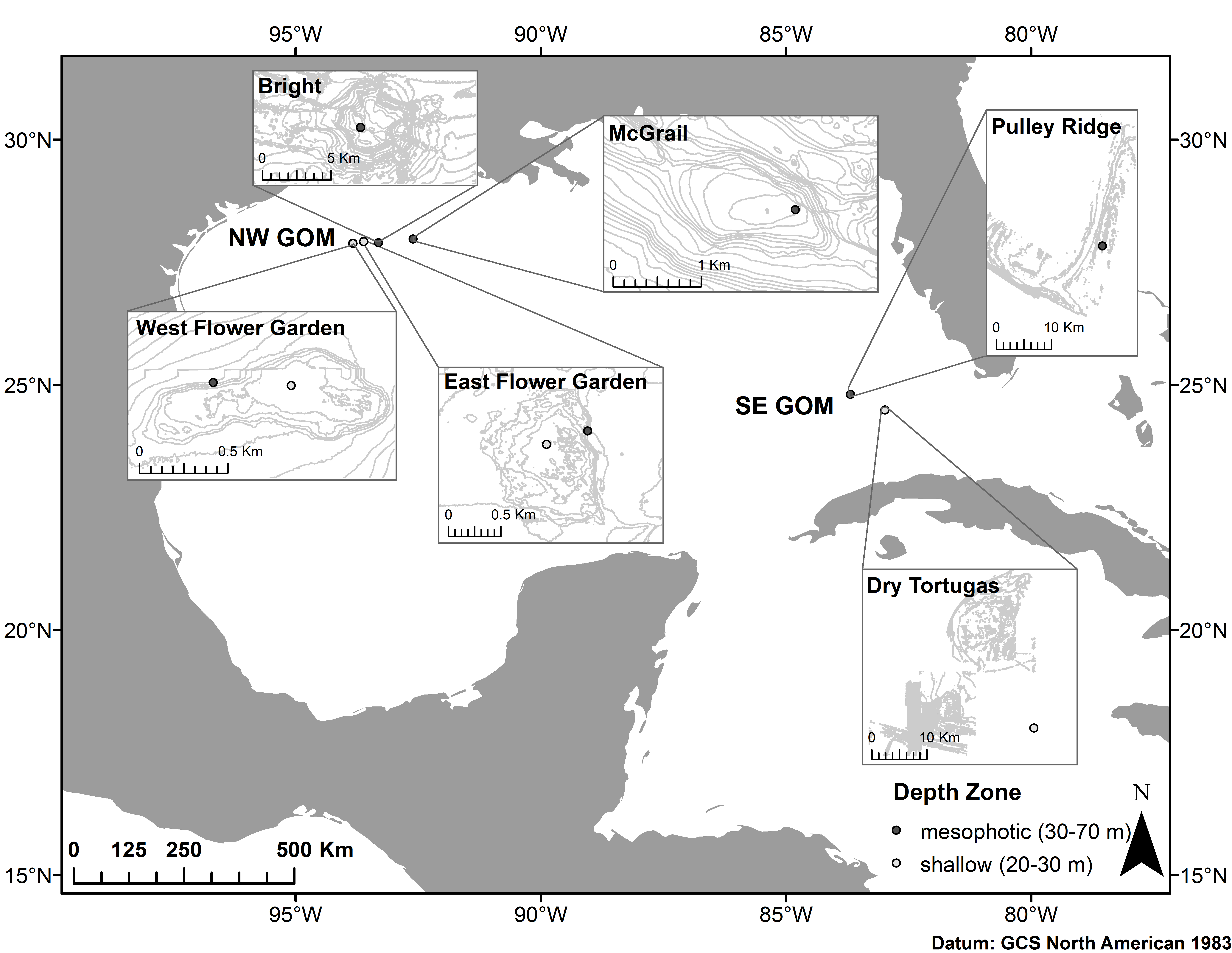

### Supplemental Fig 2

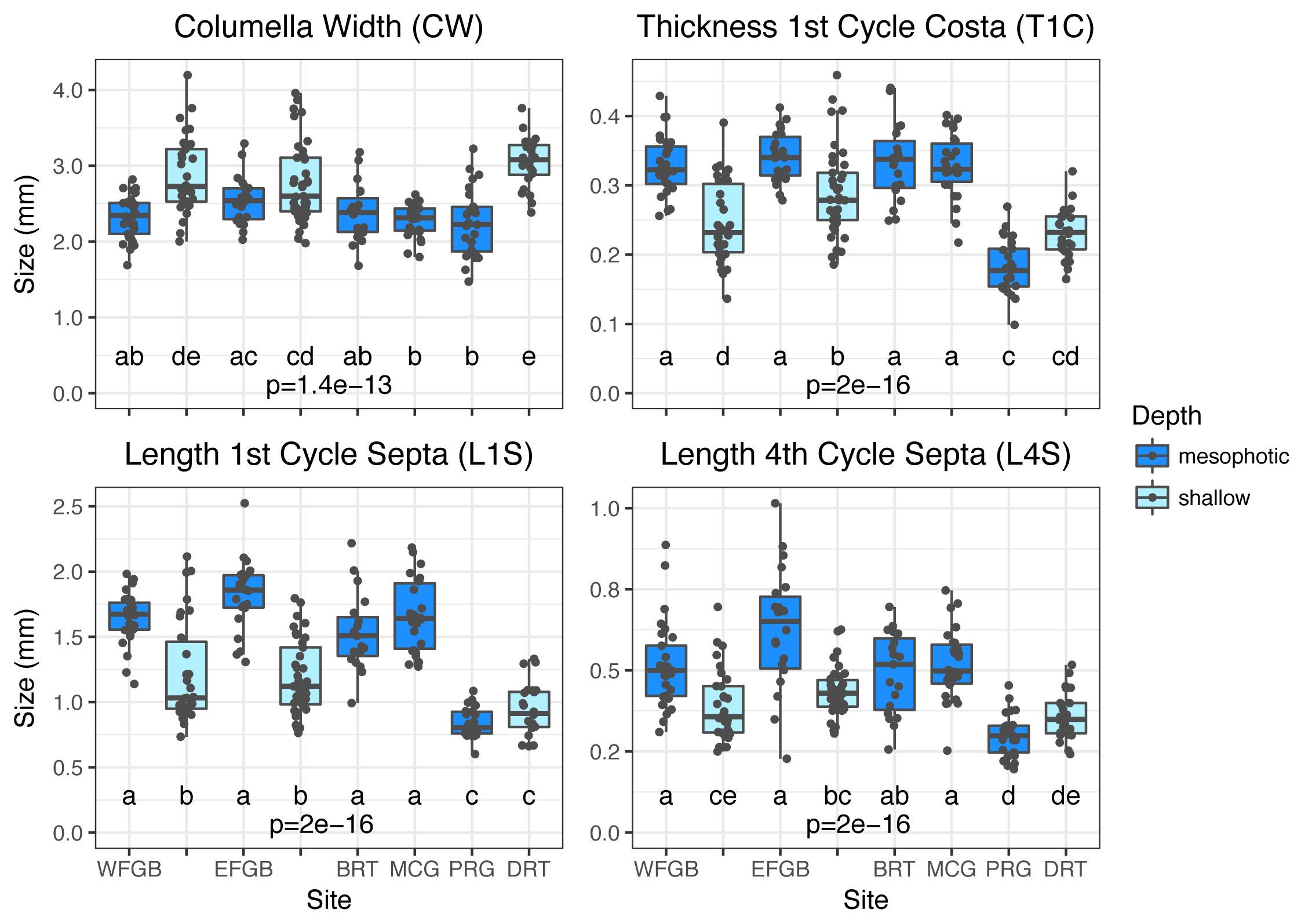

### Supplemental Fig 3

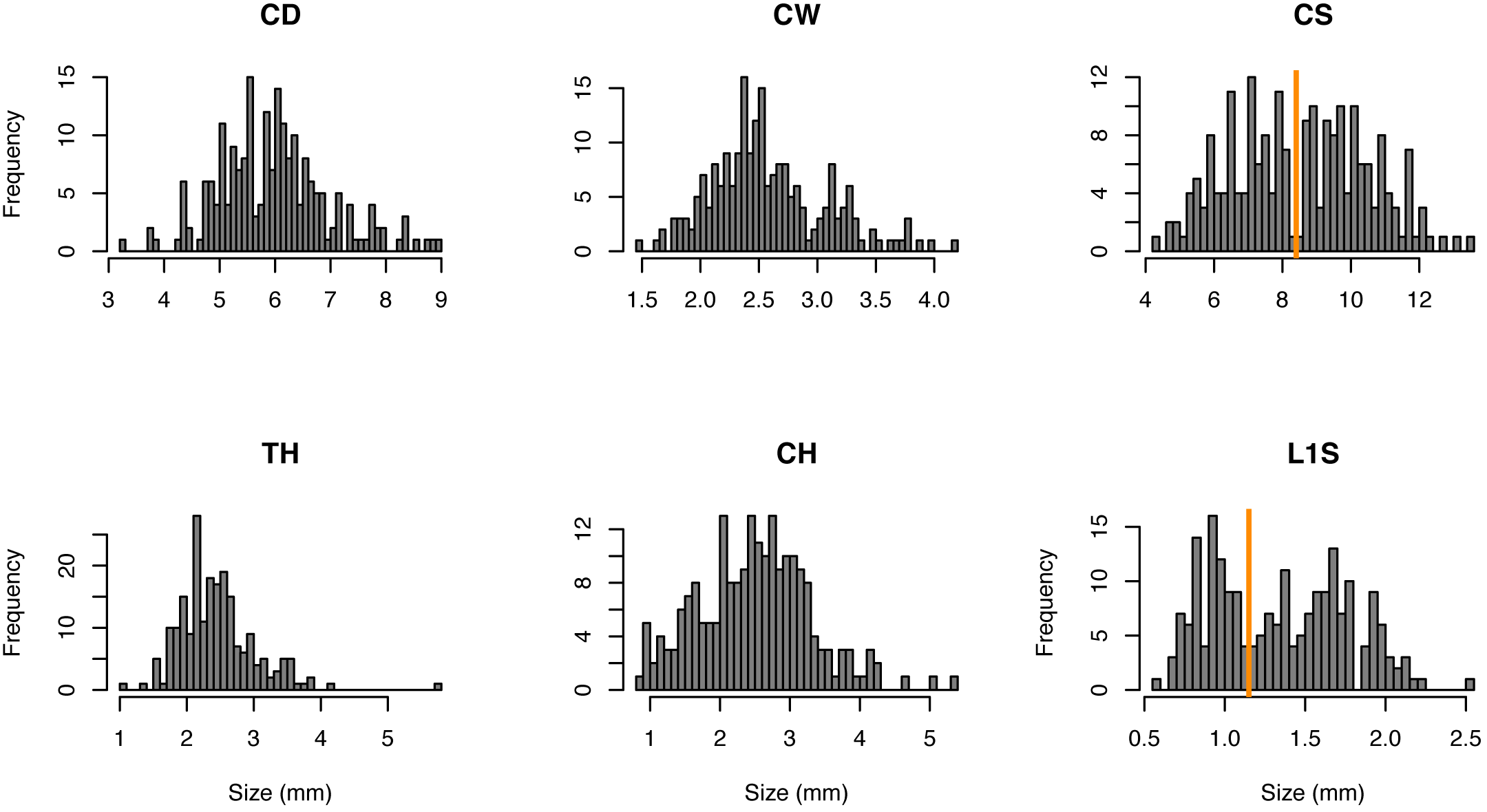

### Supplemental Fig 4

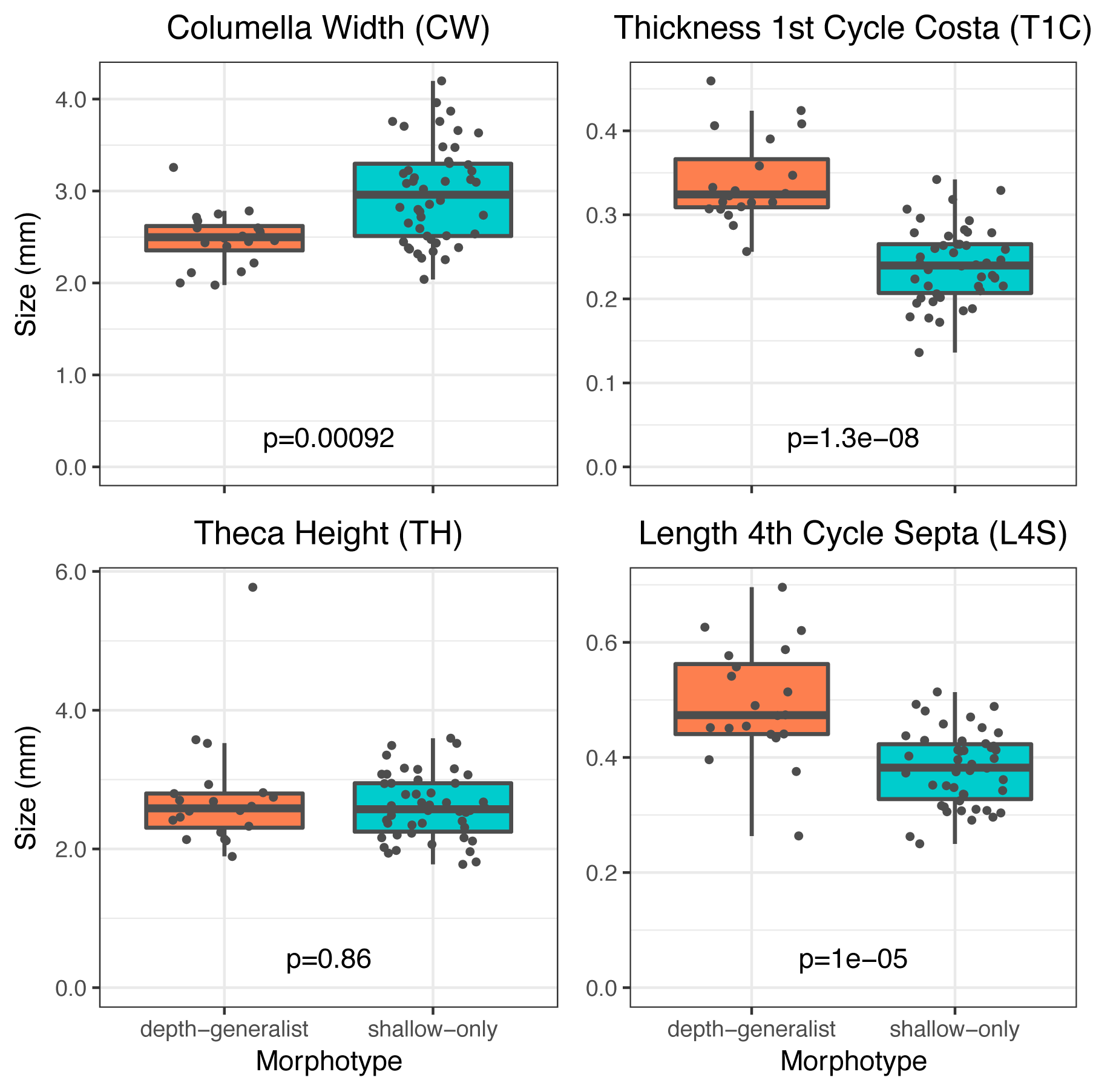

### Supplemental Fig 5

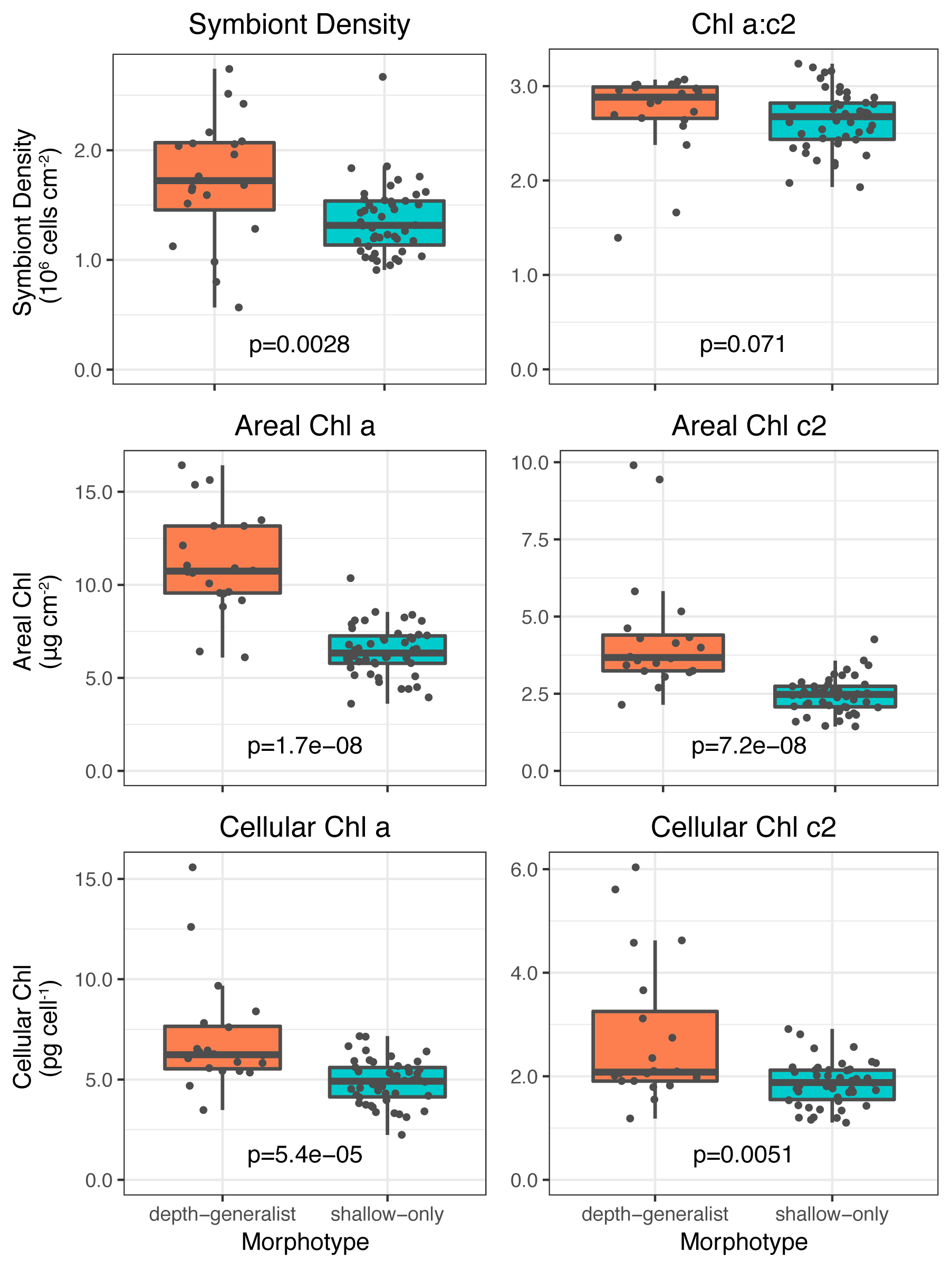

### Supplemental Fig 6

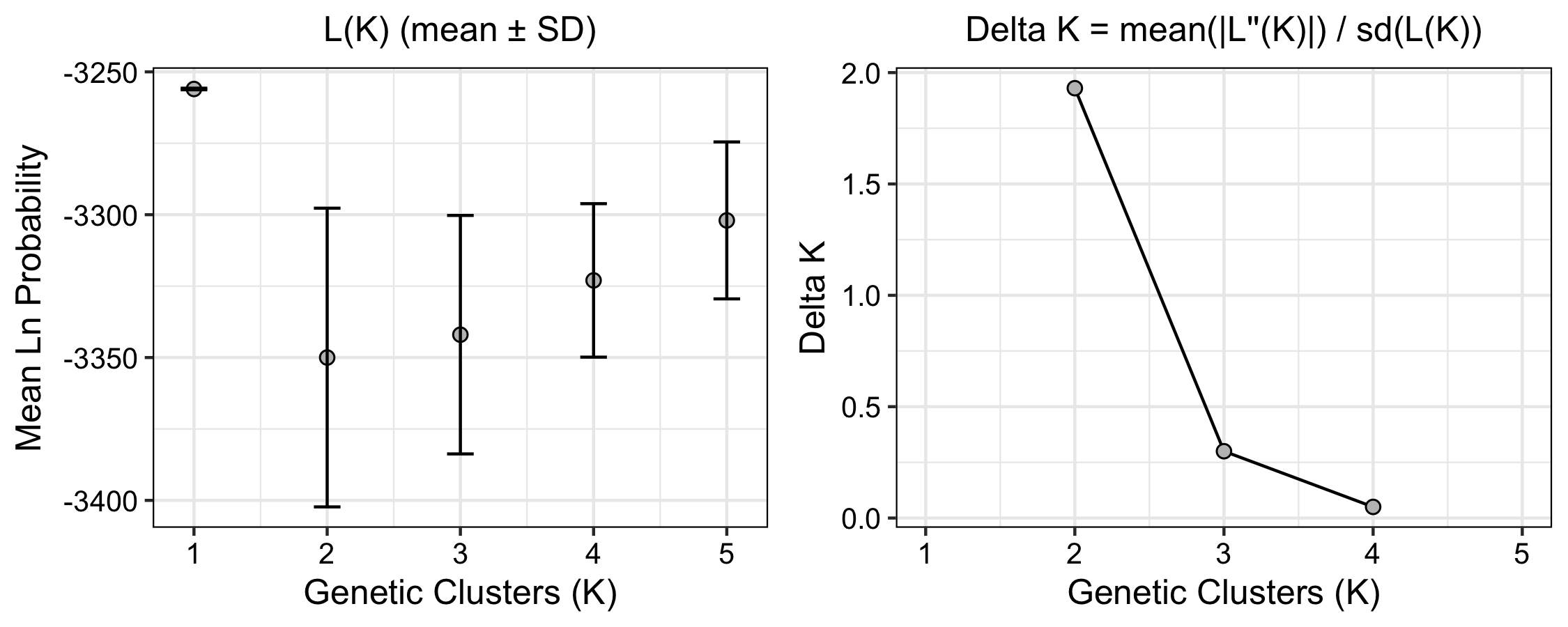
